# GenomeBits Characterization of MPXV

**DOI:** 10.1101/2022.06.21.497069

**Authors:** E. Canessa

## Abstract

Genome sequences of monkeypox virus (MPVX) causing the current outbreak are being reported from an increasing number of countries. We present a letter-to-numerical sequence study via GenomeBits signal mapping in order to characterize the evolution of the MPXV via simple statistical analysis. Histograms, empirical and theoretical cumulative distribution curves and the resulting scatter plots for the base nucleotides A and C, and their complementary base nucleotides T and G, are discussed. GenomeBits may help on the surveillance of emergent infectious diseases.

---

According to WHO, there will be more cases of monkeypox identified as surveillance expands in non-endemic countries [1]. To date, around hundred genome sequences samples have been reported from confirmed cases around the world [2–4]. These indicate a close match to exported cases from Africa found in 2017 [5]. Available information suggests that human-to-human transmission occurs among people in close physical contact with a symptomatic case. Understanding better this pathogen is still a global challenge for scientific research. A global network of permanent bioinformatics surveillance, with the capacity to rapidly deploy genomic tools and statistical studies for MPXV characterization of the evolution of lineages and mutations is still much needed.

In this brief communication, we apply the new GenomeBits statistical algorithm whose salient feature is to map the nucleotide bases *α* = *A,C,T,G* (as observed along a genome sequence) into a finite alternating (±) sum series of distributed terms of binary (0,1) indicators [6]. Genomebits allows to track and analyze viral sequences in relation to its evolution status.

This new quantitative method for the characterization of distinctive patterns of complete genome sequences of length *N* considers

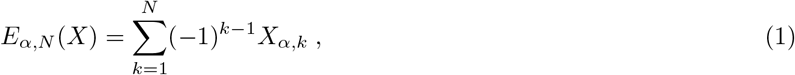

where the independently distributed terms *X_k,α_* are associated with (0,1) according to their position along the sequences, satisfying the following relation *X*_*α,k*=*N*_ = |*E*_*α,N*_(*X*) – *E*_*α,N*–1_(*X*)|. The arithmetic progression carries positive and negative signs (−1)^*k*−1^ depending on the nucleotide base position *k*.

This mapping into four binary projections of genome sequences follows previous studies on the three-base periodicity characteristic of protein-coding DNA sequences [7]. However, as a principal difference with other binary representations (see also, *e.g*., Refs [8, 9]), in Eq (1) the terms in the sums change sign. If a term *X_α,k_* in the sum series is positive at a given *k*, then the next *X*_*α,k*+1_ term is negative and vice versa. The binary indicator for the sequences is chosen sequentially starting with +1. This is illustrated in Table I, where a mapping example for converting the brief genome fragment TTTTAGTACATTAAT (consisting of 15 nucleotides) into the alternating binary array via Eq (1) is given. As illustrated in the Table, the positive and negative terms in the sums of Eq.(1) partly cancel out, allowing the sum series to diverge from zero rapidly and to become a non-Cauchy sequence type [6].

**TABLE I:**
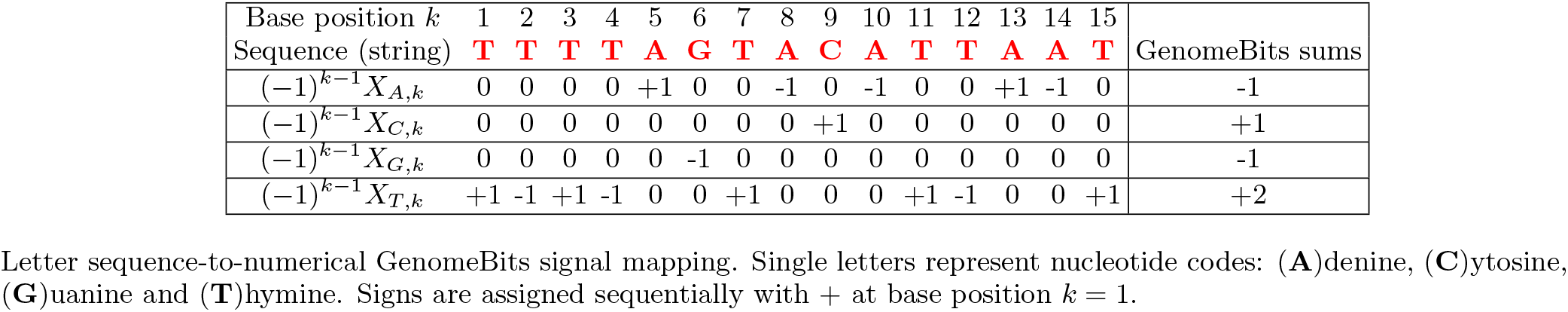
Genome sequence-*to*-GenomeBits mapping example via Eq (1) for *N* = 15.

There is a user-friendly Graphics User Interface (GUI) for the GenomeBits signal analysis method of genome sequences in FASTA format. The GUI can be downloaded freely from Github [10]. It runs under Linux Ubuntu OS. For large genomic data, say, of the order of *N* = 197204 nucleotide bases as for example for ON563414.3 monkeypox virus isolate MPXV_USA_2022_MA001 complete genome [3], the one used in the present study, the GUI requires little processing time and discards uncompleted sequences containing codification errors (usually denoted with “NNNNN” letters).

We show next some interesting statistical imprints of the intrinsic gene organization at the level of nucleotides from the above formula. In Fig 1, the results obtained via Eq.(1) for the binary sequences of each A, C, G and T nucleotide of the MPXV reported from the USA are shown. For comparison, the curves on the top display results for nucleotides A,T of the strand and on the bottom the nucleotides C,G (complementary to those of the opposite strand according to the pairing rules A-T and C-G of DNA). It is interesting to note how in the figure there are regions where the curves seems to mirror each other. This peculiar behaviour reveals coding regions of correspondence between single nucleotide structures. Also interesting is the fact that for *α* = (*A*) the total numbers of zeros is 66.4% of the whole sequence and the total number of ones is 33.6%. For *α* =(*C*) these correspond to 83.5% and 16.5%, respectively. For *α* = (*G*) one finds 83.5% and 16.5% and, similarly, for *α* = (*T*) the values become 66.6% and 33.4%.

**FIG. 1:**
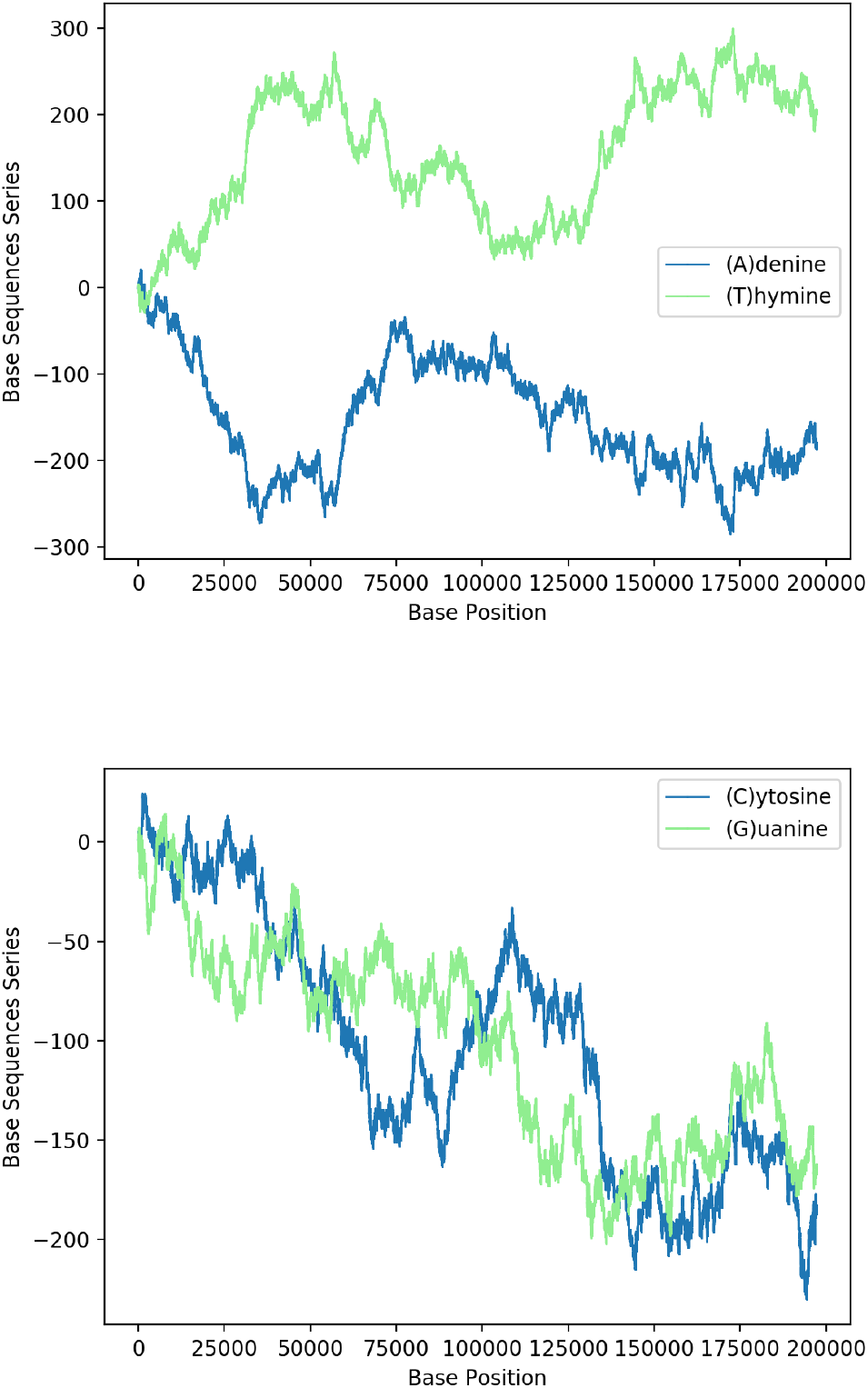
GenomeBits imprints of the genome sequence monkeypox virus isolate United States MPXV_USA_2022_MA001 (GenBank ID ON563414.3).

A first step in assessing a probability distribution of datasets is usually obtained by computing a basic histogram (*i.e*., counting the number of data points in a given bin). A standard cumulative, normalized histogram as a step function in order to visualize the empirical cumulative distribution function (CDF) for a sample is also useful [11]. In this case, one gets the probability of, and observation from, the sample not exceeding a particular x-value at a fixed y-value. However, our goal here is to compare two different datasets or alternatively two different histograms of equally sized bins.

To achieve this, a pair of histograms above and right of an scatter plot of such two datasets can help to visualize and highlight correlations at certain coloured contact points. Scatter plots with marginal distributions of this kind can be easily done using Python programming through the built-in Matplotlib’s routines: scatter(x,y), hist(x), gridspec and invert_xaxis (see, *e.g*., Ref. [12]). In Figs.2 and 3, we illustrate the (blue and green bar-graph) histograms, the empirical (red full line) and theoretical (red dotted lines) CDF curves and the resulting scatter plots for the nucleotides A and its complementary T, and C and its complementary G plotted in Fig.1. We have used number *bins* = 50 and the fits (blue line) have been calculated via expected Gaussian distribution.

**FIG. 2:**
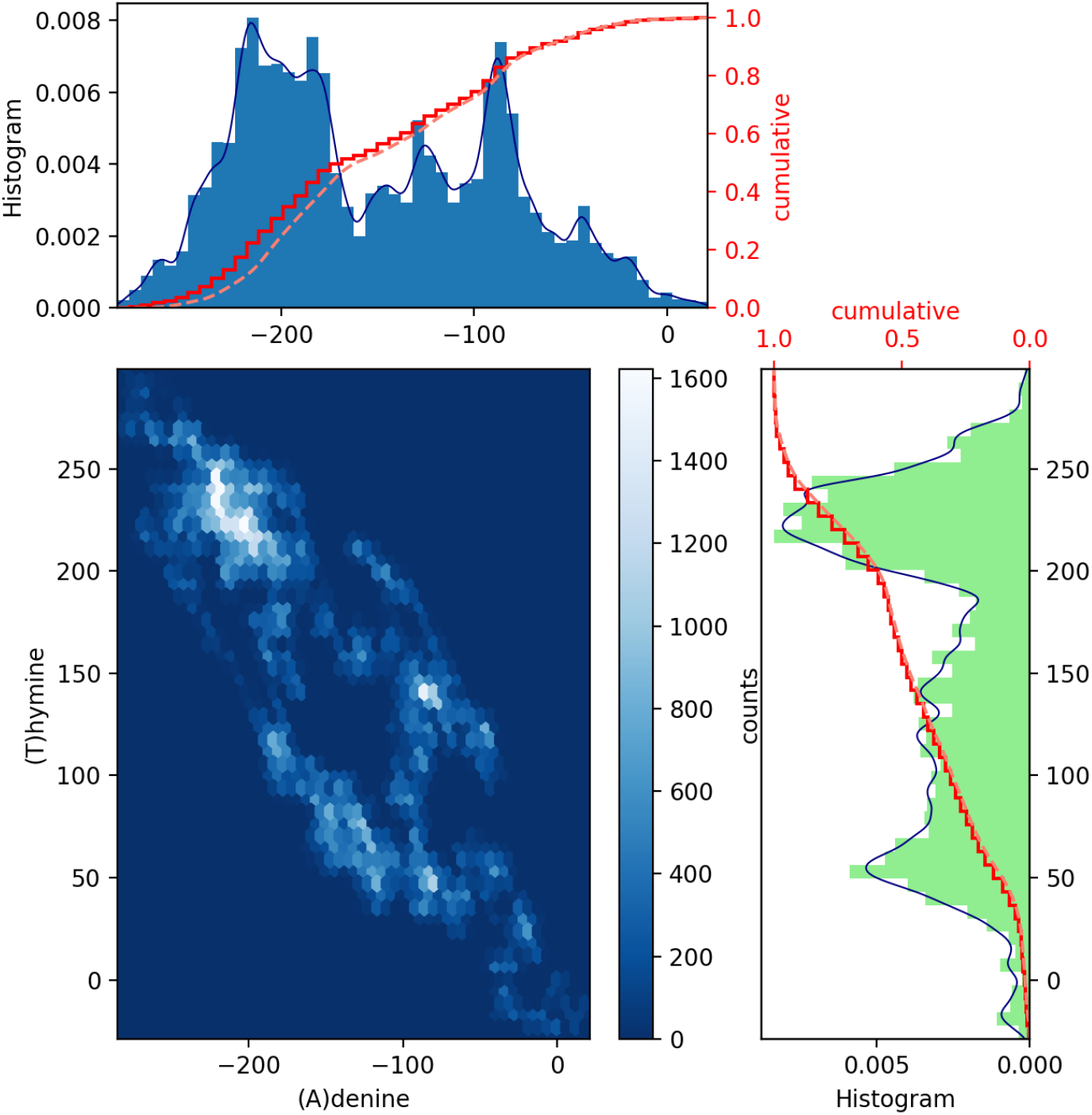
Scatter plot of GenomeBits (A)denine and (T)hymine curves with histograms for monkeypox virus (GenBank ID ON563414.3).

**FIG. 3:**
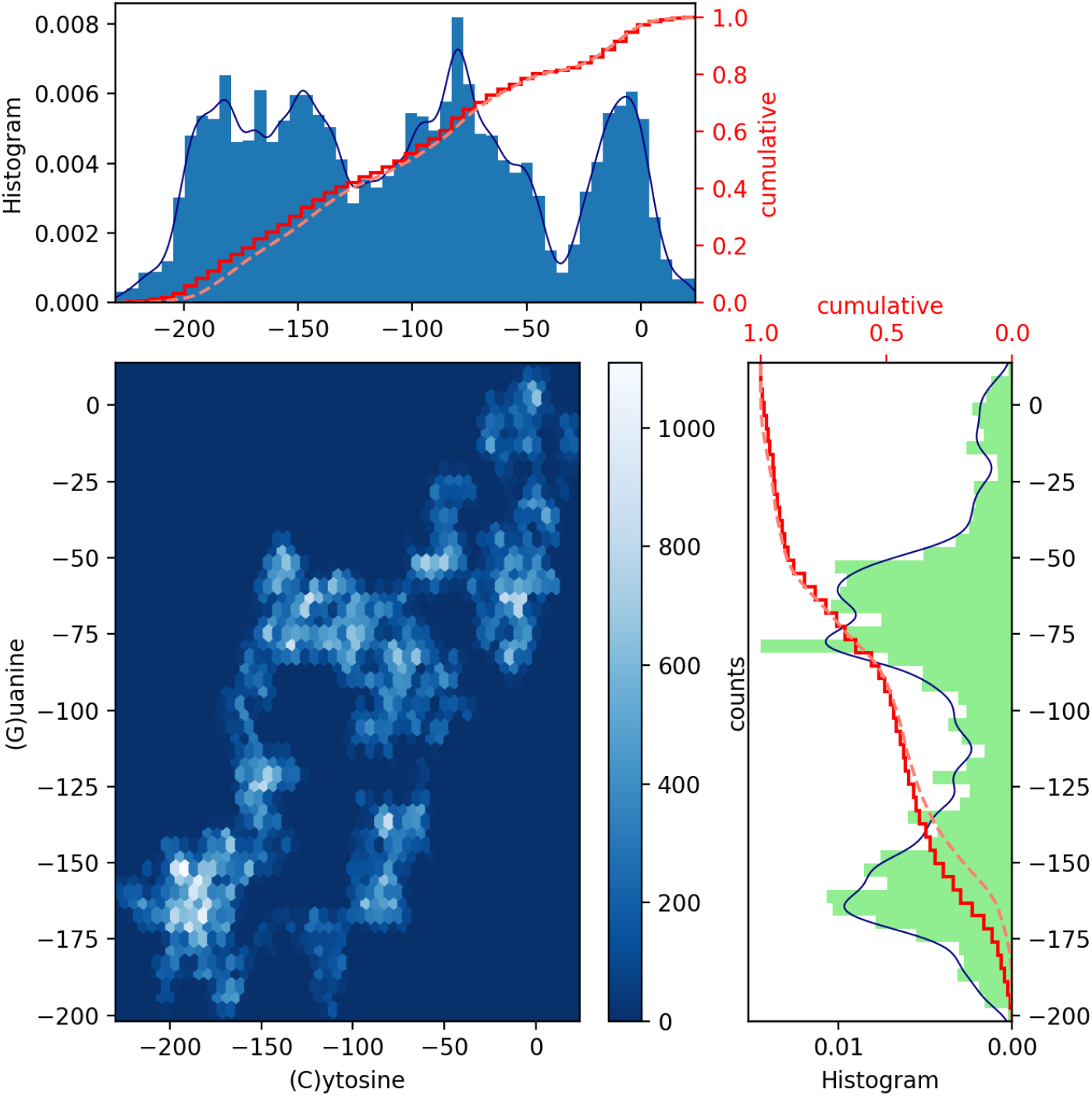
Scatter plot of GenomeBits (C)ytosine and (G)uanine curves with histograms for monkeypox virus (GenBank ID ON563414.3).

In Fig.2 and 3 we drawn scatter diagrams in an attempt to obtain correlations between GenomeBits sequences of complementary nucleotide bases. The clearer colours of the hexagon markers add a new dimension to these plots. These show how the mapped binary data is stratified along the sequences and which areas show somehow correlated points which fall along complex, rounded shapes. The markers are unequally distributed in the scatter plot and far from forming a straight line. The patterns found which summarize the underlying data do not occur by random chance.

From these results, we believe GenomeBits may help to shed light on the surveillance behind infectious diseases. By GenomeBits, we have also uncovered distinctive signals for the intrinsic gene insights in the coronavirus genome sequences for past variants of concern: Alpha, Beta and Gamma, and the variants of interest: Epsilon and Eta [6]. An study of GenomeBits into the most recent delta and omicron variants of the coronavirus pathogen can be found in Ref. [13]. The GenomeBits method focus on single nucleotide structures and, as proposed in this letter, may also provide useful information to study the evolution of the MPXV outbreak via simple statistical analysis.

